# HipSAFE: automating hip fracture detection on ultrasound imaging using deep learning

**DOI:** 10.64898/2026.03.12.711420

**Authors:** Nicholas J. Yee, Yohannes Soenjaya, Noah Kates Rose, Angela Atinga, Christine Démoré, Mansur Halai, Cari Whyne, Michael Hardisty

## Abstract

Falls among older adults can result in hip fractures that requires x-ray based assessment at emergency department (ED). Only 25.7% of patients presenting to EDs are diagnosed with a hip fracture, as such improved diagnosis prior to transportation to hospital could result in fewer hospital visits and improved triaging. Patient with hip fracture could be immediately directed to centres with orthopaedic surgeons, allowing for reduced time-to-surgery, particularly in rural communities. Ultrasound (US) imaging is portable and can identify fractures but requires expertise, particularly related to image interpretation. Deep learning may reduce operator dependence by automating image interpretation. This study aims to develop HipSAFE, a hip fracture detection tool on US, to support triaging by nurses and paramedics. We hypothesize that diagnostic accuracy will be comparable to pelvic x-ray diagnostic performance in a preclinical study. Bilateral hind limbs of 15 porcine cadavers were imaged by US-naïve operators before and after an iatrogenic hip fracture. The limbs were divided into training, validation, and test (8 femurs) sets. The training data were augmented (geometric and photometric transformations). The models included MobileNetV3 (S/L), EfficientNet-Lite (0–2), and ResNet (18/50). Using a moving average aggregation on the operator cine clips, EfficientNet-Lite0 achieved the highest performance (F1=0.944 [95% CI:0.880-0.987]; sensitivity=89.5% [78.6-97.5%]; specificity = 100.0% [100.0-100.0]). The majority voting ensemble model ranked second (F1=0.932 [0.857-0.984]). Naïve operators and radiologists had lower performance (F1=0.667 [0.596-0.758] and 0.685 [0.597-0.729]). This pre-clinical study demonstrated that HipSAFE has excellent diagnostic accuracy and there may be a role for US in improving hip trauma triaging, especially for rural and resource-constrained environments.

## Introduction

Hip fractures are common orthopaedic injuries among older adults associated with significant morbidity and mortality [1]. The estimated lifetime risk is 7.3% for women and 6.2% for men [2]. Falls from standing height are a common cause for hip fractures and subsequent presentation to the emergency department (ED) for assessment [3]. Hip pain after a fall is routinely investigated with x-ray imaging, which remains the clinical standard of care for fracture detection. However, among older adults presenting to EDs with acute hip trauma, only 25.7% are diagnosed with a hip fracture, indicating potentially avoidable medical transportation and hospital visits [3]. For patients diagnosed with fractures, delays in diagnosis and further hospital transfers for definitive orthopaedic care (i.e., surgery) are associated with prolonged time-to-surgery, increased complication rates, and higher mortality [4], [5], [6], [7], [8], [9], [10], [11], [12]. These challenges are amplified in rural and remote settings, where access to imaging infrastructure and orthopaedic surgeons is limited. Therefore, an improved pre-hospital triaging protocol for hip trauma could reduce unnecessary ED visits and medical transportation while potentially expediting care for patients with hip fractures.

Point-of-care ultrasound (US) is a portable, affordable, and radiation-free imaging modality that is reliable in detecting hip fractures [13], [14], [15]. Its affordability and portability make it particularly attractive for pre-hospital and resource-limited environments. However, the diagnostic accuracy of US is highly operator dependent, requiring sonography training that is not routinely available among first-line healthcare providers such as paramedics and nursing staff. Capturing high quality US images requires technical expertise in managing device settings and dexterity with US probe positioning to address anisotropy and limited acoustic windows. Furthermore, image interpretation of subtle sonographic fracture features is challenging and affects subsequent image collection paths. These factors raise barriers to widespread clinical adoption for using US imaging in hip trauma assessment.

Recent advances in deep learning and computer vision offer a pathway to mitigate these limitations by automating US image interpretation [16], [17]. These complex statistical models can learn hierarchical representations of fracture-related features directly from imaging data, potentially reducing reliance on expert sonographers and enabling consistent diagnostic performance among US operators without extensive sonography training [18]. A deep learning image interpretation system for US hip fracture assessment, particularly lightweight models designed for embedded systems such as mobile or point-of-care devices, may offer a real-time clinician decision support tool that would present the opportunity to have diagnostic imaging in communities with paramedics and nurses that care for older adults. To our knowledge, there are currently no published studies evaluating deep learning–based hip fracture detection models applied to ultrasound imaging.

In this preclinical study, we introduce HipSAFE, a Hip Sonographic Acute Fracture Evaluation tool, as a deep learning pipeline for automated hip fracture detection on ultrasound imaging using porcine cadaver models. We evaluated fracture classification at two levels: (1) individual image frame detection: useful for guiding subsequent image collection and (2) cine clip detection: providing a single recommendation for clinical decision making. Model performance was compared against naïve US operators, representing potential end-user paramedics and nurses who typically lack sonography training, and expert radiologists, to contextualize algorithmic performance for less frequently used imaging modality to assess hip trauma. The hypothesis is that the diagnostic accuracy will be comparable to standard pelvis x-ray performance. This study will demonstrate the potential of deep learning–assisted US imaging as a decision support triaging tool for hip fractures.

## Methodology

This preclinical study evaluated a deep learning–based pipeline for automated hip fracture detection on US imaging using porcine cadaver models. Bilateral femurs from porcine specimens were imaged before and after inducing an iatrogenic hip fracture for an intact and fractured hip dataset. No ethics approval at our institution was required for the cadaveric study as the specimens were obtained following sacrifice of the animals for secondary use.

Convolutional neural networks (CNNs) were evaluated on detecting hip fractures on US frames and clips (frames in a time series) collected by researchers unfamiliar with musculoskeletal sonography to approximate sonography collection of nurses and paramedics without specific US musculoskeletal training. The interpretations by the algorithms were compared to assessments by naïve operators as well as diagnostic radiologists.

### Data collection

Prospective US data were collected from the bilateral hind limbs of 15 juvenile porcine cadavers (30 hips) using a Philips EPIQ 7G Ultrasound System (Koninklijke Philips N.V., Amsterdam, Netherlands). Imaging was performed with linear probe with a frequency range of 5.0-12.0 MHz. US B-mode imaging were acquired in DICOM format and converted to MP4 cine clips for downstream processing and model training.

Pertrochanteric hip fractures were induced through a posteromedial approach using a bone saw and osteotome with care to avoid introducing a tissue-air interface on the anterolateral imaging plane. Direct palpation of a bicortical break confirmed a complete fracture in all hips. All cine clips were captured after the dissection to reduce the risk of the DL models learning spurious signals from the dissection. Cine clips were captured before and after the iatrogenic fracture resulting in intact and fractured datasets for each femur.

The datasets were split at the femur level into training (16 femurs), validation (6 femurs), and held-out test (8 femurs) datasets to prevent data leakage. Two types of datasets were collected on the same femurs: anatomical dataset and operator cine clip dataset. For both datasets, the fractured limb was manipulated with each imaging session to increase variability in fracture positioning.

For the anatomical dataset, the cine clips were acquired from the femoral head to the base of the lesser trochanter and along the subtrochanteric diaphysis. Standardized probe orientations were used including longitudinal, transverse, and oblique views to capture parallel, perpendicular, and oblique views of the bone. The training and validation datasets included 49,076 image frames (11,834 fractured bone; 23,995 intact bone; 13,247 tissue). The test dataset contained 14,958 images (4,123 fractured bone; 6,677 intact bone; 4,158 tissue). A balanced distribution of longitudinal, oblique, and transverse views was collected with an emphasis on the intertrochanteric region. (Table 1).

**Table 1:** Distribution of image frames of different anatomical regions imaged for the training/validation and test datasets.

|  | Train + validation | Test |
| --- | --- | --- |
| Longitudinal intertrochanter | 7918 (22%) | 2533 (23%) |
| Oblique intertrochanter | 8041 (22%) | 2497 (23%) |
| Transverse intertrochanter | 7674 (21%) | 2500 (23%) |
| Longitudinal shaft | 4164 (11%) | 1113 (10%) |
| Oblique shaft | 4077 (11%) | 1140 (10%) |
| Transverse shaft | 3955 (11%) | 1017 (9%) |

The operator cine clip dataset consisted of 78 test-set femur cine clips collected by 31 US-naïve operators. The operators had no prior sonographic training. Before the imaging session, they were shown images of intact and fractured bones on US along the different anatomical planes and orientated to the femur position on the cadaver. They were blinded on the fracture status, and they received no feedback between sessions until the study completion to minimize a learning effect.

Diagnostic radiologists (a board-certified musculoskeletal radiologist and a senior radiology resident) read the cine clips collected by the operators. They had no prior experience in reading porcine US imaging. Operators made assessments of the fracture status during US acquisition. The operators and diagnostic radiologists indicated the presence or absence of a fracture.

### Data preprocessing

All frames were cropped to retain only the ultrasound field of view and exclude overlays. Data augmentation was applied only to the training set and included rotation, translation, scaling, horizontal flipping, resolution perturbation, perspective transformation, colour jitter, and Gaussian blurring (Table 2 and Figure 1).

**Table 2:** Data augmentation hyperparameters applied during model training to improve generalization and reduce overfitting.

| Augmentation | Category | Details | Application probability |
| --- | --- | --- | --- |
| Horizontal flip | Geometric | Reflected across the y-axis | 0.5 |
| Random affine transformation | Geometric | Rotation $\pm 20^\circ$; translation $\pm 20\%$; scale 0.8–1.2 | 0.25 |
| Resolution perturbation | Geometric | Downsampled to $256 \times 256$ then upsampled to $640 \times 640$ (bilinear interpolation) | 0.25 |
| Perspective transformation | Geometric | Distortion scale = 0.4 | 0.25 |
| Sharpness adjustment | Photometric | Sharpness factor = 2.0 | 0.25 |
| Colour jitter | Photometric | Brightness, contrast, saturation = $\pm 0.2$ | 0.25 |
| Gaussian blur | Photometric | Kernel size = 3; $\sigma = 0.1$–2.0 | 0.25 |

**Figure 1.**
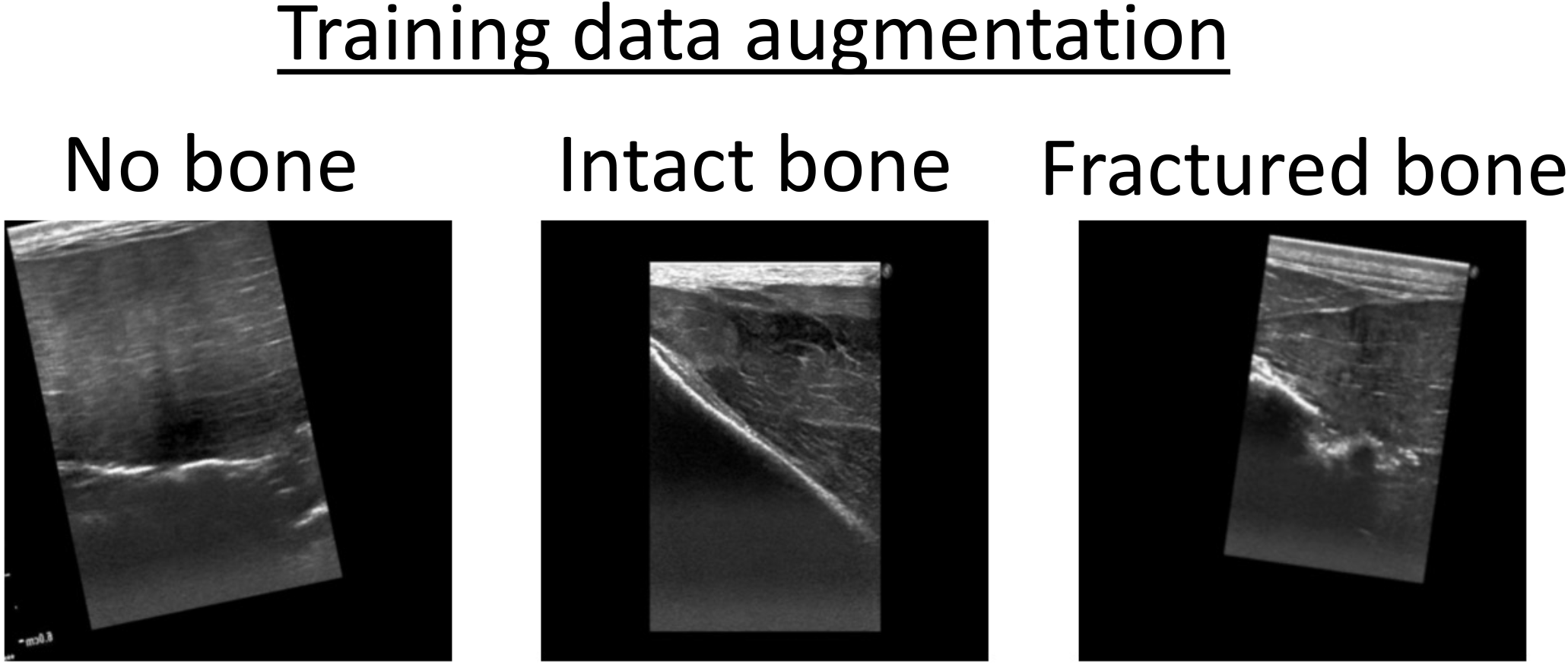
Example data augmentations on US images of A) soft tissue/no bone; B) intact bone; C) fractured bone.

### Model selection

For the frame-level feature extractor models, both standard and mobile-friendly deep learning architectures were evaluated. Standard vision models included ResNet-18 and ResNet-50 [19], while mobile-optimized architectures included MobileNetV3 (Small and Large) [20] and EfficientNet-Lite (0, 1, and 2) [21]. Mobile-friendly models were included to assess architectures designed for reduced parameter count and computational complexity through depthwise separable convolutions and compound scaling, enabling efficient feature extraction on embedded devices, which could enable direct inference on point-of-care US machines. All models were pre-trained on ImageNet to leverage transfer learning for improved model convergence on our limited medical imaging data [22].

The deep learning models were composed of sequential convolutional blocks with batch normalization and non-linear activation functions, followed by global pooling layers for spatial feature aggregation before passing to a multilayer perceptron for classification. ResNet architectures incorporated identity-based residual connections to minimize vanishing and exploding gradients in deeper networks. MobileNet architectures employed depthwise separable convolutions and inverted residual bottleneck blocks to reduce computational complexity and parameter count. EfficientNet-Lite architectures used compound scaling of network depth, width, and input resolution. For all models, the classification head with He-initialization consisted of a fully connected layer with 3 output nodes corresponding to no bone present, intact bone, and fractured bone [23].

The feature extractor models were trained for up to 100 epochs using cross-entropy loss with the Adam optimizer, a batch size of 32, and an initial learning rate of 0.000001. Early stopping was employed based on validation loss. A lambda learning rate scheduler was used, where the learning rate was multiplied by a factor of 0.1 starting at epoch 75. During the end-to-end training cycle, the best model weights was selected by the validation loss. All training were conducted on an Apple M2 Max chip with 32GB RAM. Model interpretability was assessed using Gradient-weighted Class Activation Mapping (Grad-CAM) to visualize class-discriminative regions contributing to the predictions [24].

### Prediction on cine clips

For fracture identification on the cine clips collected by the operators, the feature extractors produced frame-level fracture probabilities that were aggregated using a moving average (Figure 2). Cine clips were subsampled to include every 10^th^ frame. Three frames within the subsampled clip were used in the moving average. The feature extractors were calibrated to the operator cine clips using the cine clips collected on the validation femurs. A clip was classified as fractured if the maximal aggregated fracture probability exceeded the threshold determined by maximizing the Youden index on the validation dataset.

**Figure 2.**
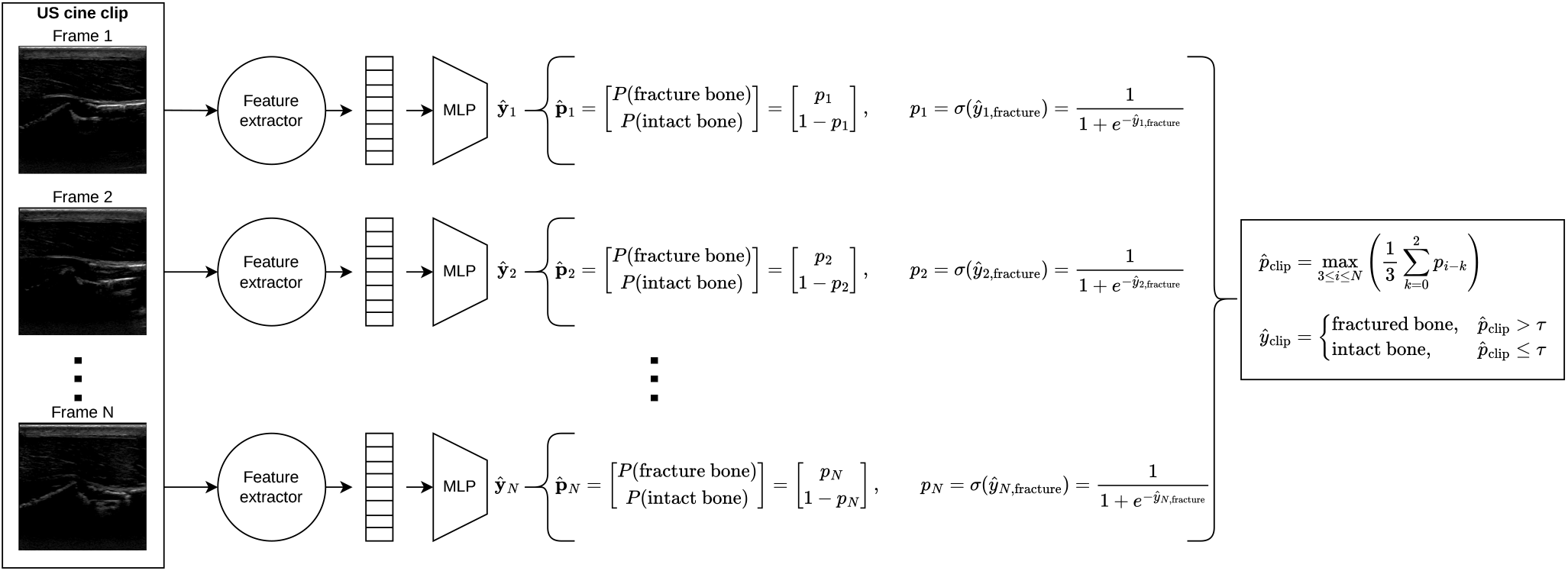
HipSAFE data pipeline for performing image frame feature extraction and cine clip hip fracture detection using the maximum fracture probability over a 3-second moving average and fracture threshold, τ.

An ensemble model was created through a majority voting of all the models’ predictions on whether the cine clip includes fractured bone.

### Metrics and statistical analysis

Model performance was evaluated at both the image frame and cine clip levels on the held-out test femurs. Metrics included accuracy, precision/positive predictive value (PPV), recall/sensitivity, specificity, F1 score, and area under the receiver operating characteristic curve (AUROC) for each model. Uncertainty estimates were obtained using bootstrapping with replacements for 1,000 iterations. Results were reported as mean values with 95% confidence intervals. Comparative analyses between deep learning models, naïve operators, and radiologists used the Friedman test followed by pairwise Wilcoxon signed-rank tests with Holm correction (α = 0.01). Interrater reliability between the two radiologists was quantified using Cohen’s kappa.

## Results

### US image frames

Among mobile-friendly architectures evaluated at the image frame level, Mobilenet V3 Small (1.5 million parameters) and EfficientNet-Lite0 (3.4 million parameters) demonstrated the highest performance, achieving an F1 score of 0.809 (0.764-0.851) and 0.774 (0.724-0.818), respectively (Table 3). Their sensitivity was 68.2% (61.9-74.3%) and 63.2% (56.7-69.2%), specificity was 99.8% (99.4-100.0%) and 100.0% (100.0-100.0%), and PPV was 99.4% (97.7-100.0%) and 100.0% (100.0-100.0%), respectively. Both models consistently outperform their larger variants (all *P*<.01). The models showed near-perfect specificity and PPV but moderate sensitivity, indicating it is highly reliable for confirming fractures while missing a meaningful proportion of fractures.

**Table 3:** Image classifier performance of mobile-friendly and standard feature extractor models on the test anatomical dataset. The bolded result was the highest for the metric. The mean and 95% confidence interval were shown.

| Model | Number of parameters (millions) | Accuracy (%) | Precision/Positive predictive value (%) | Recall/Sensitivity (%) | Specificity (%) | F1 Score | AUROC |
| --- | --- | --- | --- | --- | --- | --- | --- |
| Resnet18 | 11.2 | 88.2 (86.1-90.3) | **100.0 (100.0-100.0)** | 56.9 (50.6-63.0) | **100.0 (100.0-100.0)** | 0.725 (0.672-0.773) | 0.976 (0.964-0.986) |
| Resnet50 | 23.5 | 90.5 (88.5-92.3) | 99.4 (98.0-100.0) | 65.7 (59.6-71.8) | 99.8 (99.5-100.0) | 0.791 (0.745-0.834) | 0.986 (0.976-0.993) |
| Mobilenet V3 Small | 1.5 | **91.2 (89.2-93.0)** | 99.4 (97.7-100.0) | **68.2 (61.9-74.3)** | 99.8 (99.4-100.0) | **0.809 (0.764-0.851)** | **0.987 (0.981-0.993)** |
| Mobilenet V3 Large | 4.2 | 86.1 (83.7-88.3) | 97.6 (94.4-100.0) | 50.6 (44.5-56.8) | 99.5 (98.9-100.0) | 0.667 (0.609-0.719) | 0.919 (0.894-0.941) |
| Efficientnet-Lite0 | 3.4 | 89.9 (87.8-91.7) | **100.0 (100.0-100.0)** | 63.2 (56.7-69.2) | **100.0 (100.0-100.0)** | 0.774 (0.724-0.818) | 0.984 (0.974-0.991) |
| Efficientnet-Lite1 | 4.1 | 88.2 (85.8-90.3) | **100.0 (100.0-100.0)** | 56.9 (49.8-63.1) | **100.0 (100.0-100.0)** | 0.725 (0.665-0.774) | 0.974 (0.964-0.982) |
| Efficientnet-Lite2 | 4.8 | 86.4 (84.2-88.5) | 98.4 (95.6-100.0) | 51.0 (45.1-57.1) | 99.7 (99.2-100.0) | 0.672 (0.615-0.723) | 0.969 (0.954-0.982) |

Among standard vision models, ResNet-50 (23.5 million parameters) achieved the highest performance, achieving an F1 score of 0.791 (0.745-0.834), sensitivity of 65.7% (59.6-71.8%), specificity of 99.8% (99.5-100.0%), and PPV of 99.4% (98.0-100.0%). It had comparable performance to Mobilenet V3 Small and EfficientNet-Lite0 despite being substantially larger model (*P*=.41 and .45, respectively)

Grad-CAM demonstrated that the models localized the intact and fractured bone well while having an ambiguous region attribution for surrounding soft tissue (Figure 3). Comparable inference patterns and attribution maps on the original and horizontal flipped image suggests reflection invariance properties (Figure 4). The model consistently highlighted clinically relevant cortical regions in both axial and longitudinal views. However, the attribution maps for the fracture classification mistakenly includes the growth plate, which is the bone-cartilage interphase where bone growth occurs (Figure 4). Other failure modes identified, including the misclassification of intermuscular fascia as bone, indicating that while salient anatomical features were used for prediction, spurious features were contributing to the feature extractor model predictions.

**Figure 3.**
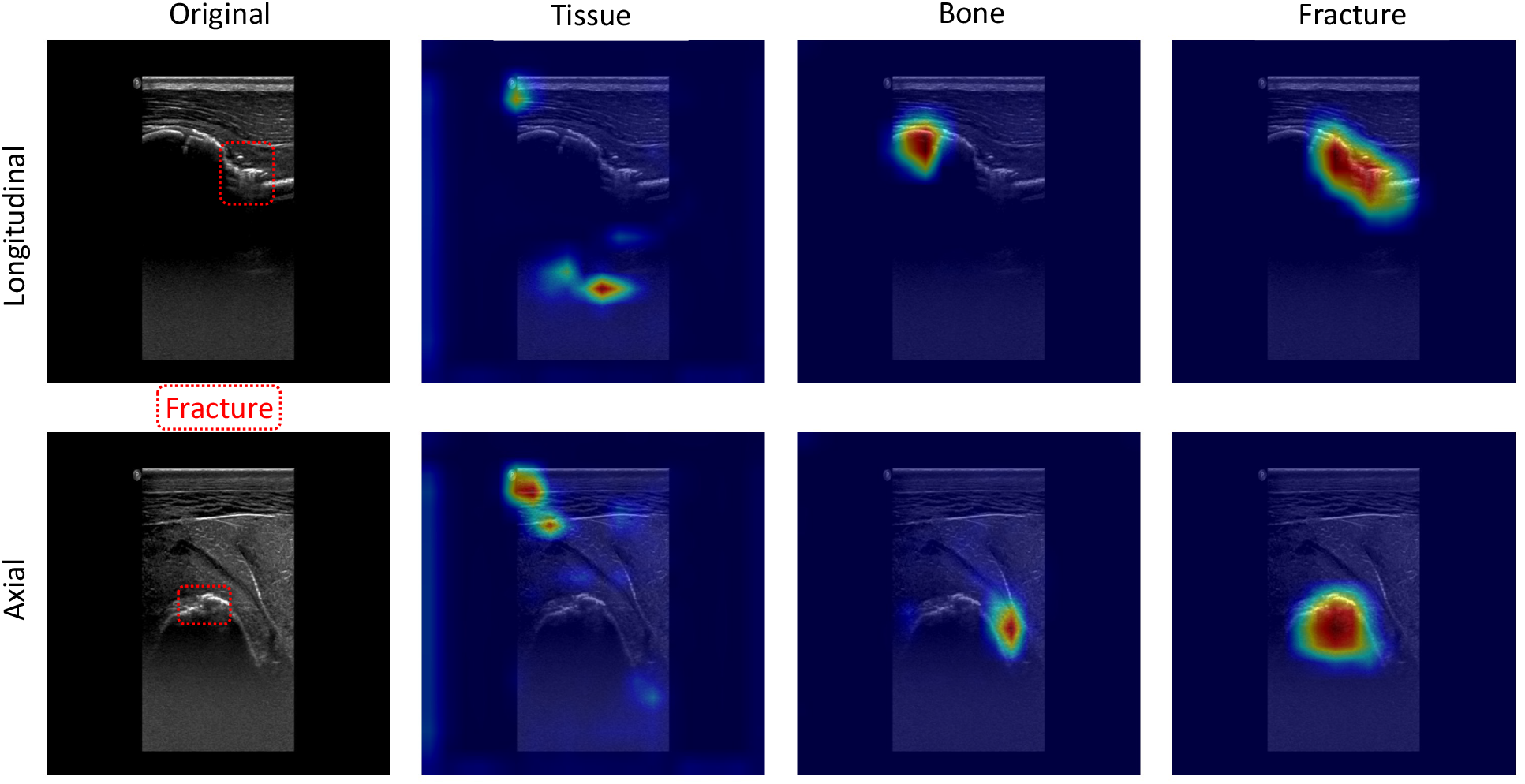
Grad-CAM localizations of intact and fractured bone along longitudinal (top) and axial (bottom) US images of the bone.

**Figure 4.**
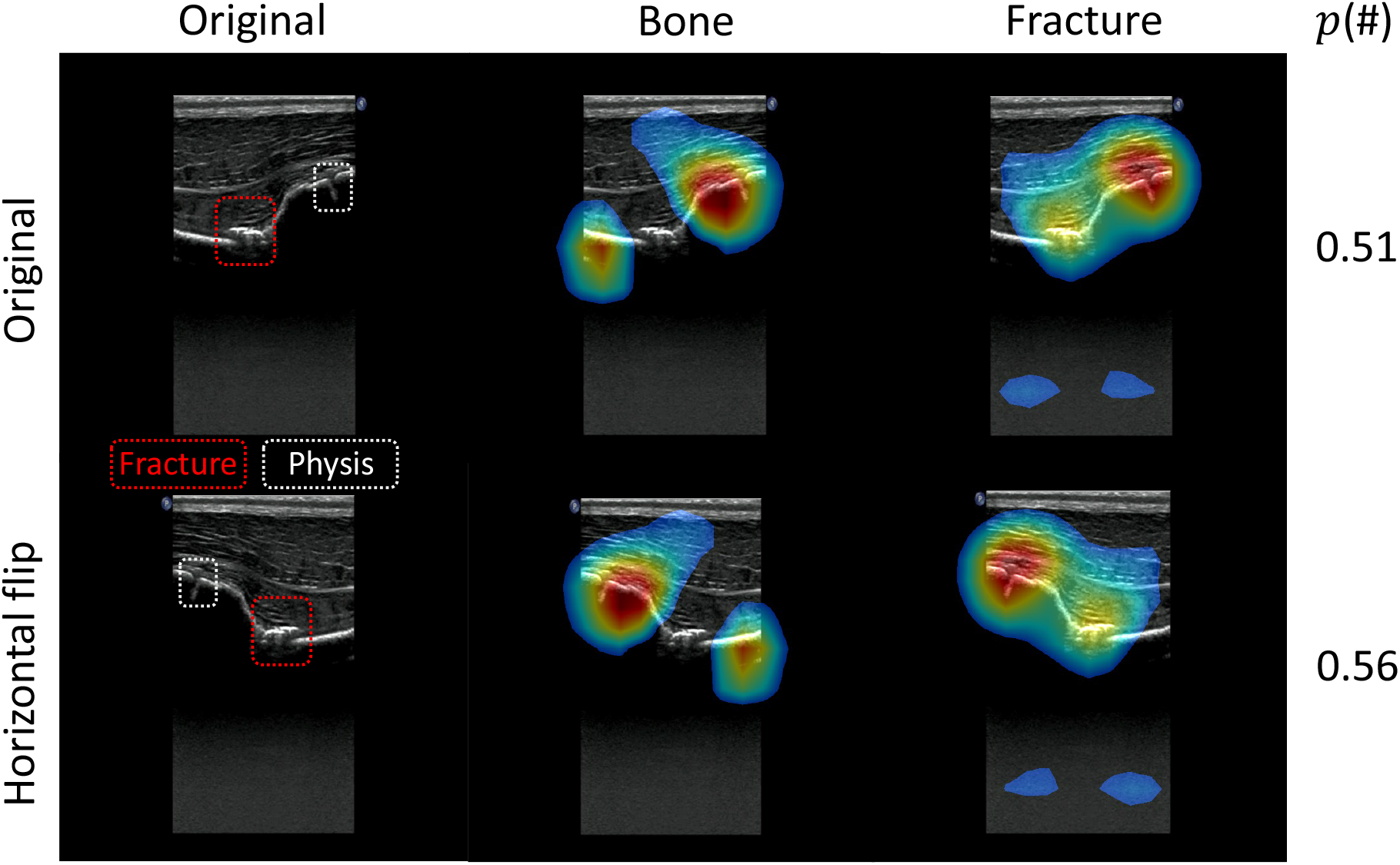
**Reflection invariance demonstrated visually through Grad-CAM attribution maps of the same image that has been reflected along the vertical axis. Region with low contribution to the fracture class were removed to highlight the attribution pattern. p(#) indicate the predicted fracture pattern. Fracture pattern included the fracture and the growth plate (intact bone).**

### Cine clips

The threshold moving average aggregation method with the different feature extraction models was used on the operator cine clips (Table 4). EfficientNet-Lite0 achieved an excellent overall performance with a F1 score of 0.944 (0.880-0.987), sensitivity of 89.5% (78.6-97.5%), specificity of 100.0% (100.0-100.0%), and PPV of 100.0% (100.0-100.0%). It was significantly better than EfficientNet-Lite2 (*P*<.01). No other models were significantly different in performance.

**Table 4:** Fracture identification on cine clips of test set porcine cadaveric hips collected by naïve US operators and read by them, the radiologists, and deep learning models. The bolded result was the highest for the metric. The mean and 95% confidence interval were shown.

| Model | Accuracy (%) | Precision/Positive predictive value (%) | Recall/Sensitivity (%) | Specificity (%) | F1 Score | AUROC |
| --- | --- | --- | --- | --- | --- | --- |
| Naïve operators | 57.7 (50.3 - 69.3) | 54.1 (44.8 - 66.9) | 86.8 (76.4 - 95.6) | 30.0 (21.7 - 47.5) | 0.667 (0.596 - 0.758) | N/A |
| Avg radiologists | 70.5 (61.2 - 74.1) | 71.4 (58.4 - 77.1) | 65.8 (55.7 - 74.3) | 75.0 (61.4 - 79.0) | 0.685 (0.597 - 0.729) | N/A |
| Resnet18 | 88.5 (80.8-94.9) | 83.7 (72.3-93.9) | **94.7 (86.5-100.0)** | 82.5 (70.4-93.5) | 0.889 (0.810-0.955) | 0.966 (0.923-0.995) |
| Resnet50 | 89.7 (83.3-96.2) | 87.5 (76.7-96.9) | 92.1 (82.6-100.0) | 87.5 (77.1-96.8) | 0.897 (0.816-0.958) | **0.982 (0.957-0.999)** |
| Mobilenet V3 Small | 76.9 (67.9-84.6) | 70.0 (56.9-81.3) | 92.1 (81.6-100.0) | 62.5 (46.2-76.1) | 0.795 (0.693-0.871) | 0.895 (0.822-0.955) |
| Mobilenet V3 Large | 84.6 (76.9-92.3) | 93.3 (82.9-100.0) | 73.7 (59.6-87.5) | 95.0 (87.1-100.0) | 0.824 (0.712-0.912) | 0.916 (0.841-0.978) |
| Efficientnet-Lite0 | **94.9 (89.7-98.7)** | **100.0 (100.0-100.0)** | 89.5 (78.6-97.5) | **100.0 (100.0-100.0)** | **0.944 (0.880-0.987)** | 0.978 (0.945-0.997) |
| Efficientnet-Lite1 | 87.2 (78.2-93.6) | 86.8 (74.3-97.1) | 86.8 (74.4-97.0) | 87.5 (75.6-97.4) | 0.868 (0.773-0.938) | 0.936 (0.863-0.991) |
| Efficientnet-Lite2 | 74.4 (64.1-83.3) | 82.1 (66.7-95.8) | 60.5 (46.1-75.0) | 87.5 (76.7-97.2) | 0.697 (0.567-0.812) | 0.823 (0.726-0.908) |
| Majority Voting Ensemble | 93.6 (87.2-98.7) | 97.1 (90.0-100.0) | 89.5 (78.6-97.5) | 97.5 (91.4-100.0) | 0.932 (0.857-0.984) | 0.935 (0.874-0.984) |

The majority voting ensemble model had a F1 score of 0.932 (0.857-0.984), sensitivity of 89.5% (78.6-97.5%), specificity of 97.5% (91.4-100.0%), and PPV of 97.1% (90.0-100.0%). It had the second highest F1 score after the EfficientNet-Lite0 model.

The naïve operators and average radiologists achieved a F1 score of 0.667 (0.596 - 0.758) and 0.685 (0.597 - 0.729), sensitivity of 86.8% (76.4 - 95.6%) and 65.8% (55.7 - 74.3%), specificity of 30.0% (21.7 - 47.5%) and 75.0% (61.4 - 79.0%), and PPV of 54.1% (44.8 - 66.9%) and 71.4% (58.4 - 77.1%), respectively.

Each femur had multiple operators collect cine clips resulting in different perspectives of the same bone. On the different clips for the same femur, the deep learning models were more consistent with determining whether the bone was fractured compared to the naïve operators and radiologists (Table 5). Inter-rater agreement among the radiologists was fair(κ=0.22).

**Table 5:** Consistency in identifying fractures on the cine clips collected by naïve US operators and interpreted by naïve operators, radiologists, and deep learning models. Percentages represent the proportion of clips per femur classified as fractured.

| Femur ID | Fractured? | Number of operator clips | Naïve operators (%) | Avg radiologists (%) | Resnet18 (%) | Resnet50 (%) | Mobilenet V3 Small (%) | Mobilenet V3 Large (%) | Efficientnet-Lite0 (%) | Efficientnet-Lite1 (%) | Efficientnet-Lite2 (%) | Majority Voting Ensemble (%) |
| --- | --- | --- | --- | --- | --- | --- | --- | --- | --- | --- | --- | --- |
| 1 | No | 10 | 70 | 45 | 10 | 0 | 70 | 0 | 0 | 10 | 10 | 0 |
| 2 | No | 10 | 70 | 35 | 10 | 0 | 60 | 0 | 0 | 10 | 0 | 0 |
| 3 | No | 3 | 67 | 17 | 33 | 33 | 33 | 33 | 0 | 33 | 33 | 33 |
| 4 | No | 3 | 67 | 50 | 67 | 67 | 33 | 0 | 0 | 33 | 67 | 0 |
| 5 | No | 2 | 100 | 0 | 0 | 0 | 0 | 50 | 0 | 0 | 0 | 0 |
| 6 | No | 2 | 50 | 0 | 0 | 0 | 0 | 0 | 0 | 0 | 0 | 0 |
| 7 | No | 5 | 60 | 0 | 40 | 20 | 0 | 0 | 0 | 20 | 20 | 0 |
| 8 | No | 5 | 80 | 0 | 0 | 20 | 0 | 0 | 0 | 0 | 0 | 0 |
| 1 | Yes | 5 | 100 | 90 | 80 | 80 | 100 | 60 | 80 | 60 | 60 | 80 |
| 2 | Yes | 5 | 80 | 70 | 100 | 80 | 100 | 60 | 80 | 80 | 40 | 80 |
| 3 | Yes | 3 | 100 | 67 | 100 | 100 | 100 | 100 | 100 | 100 | 100 | 100 |
| 4 | Yes | 3 | 67 | 17 | 100 | 100 | 100 | 100 | 100 | 100 | 100 | 100 |
| 5 | Yes | 3 | 67 | 50 | 100 | 100 | 100 | 100 | 100 | 100 | 100 | 100 |
| 6 | Yes | 3 | 100 | 67 | 100 | 100 | 100 | 67 | 100 | 100 | 100 | 100 |
| 7 | Yes | 8 | 88 | 69 | 100 | 100 | 88 | 63 | 88 | 88 | 25 | 88 |
| 8 | Yes | 8 | 88 | 69 | 88 | 88 | 75 | 75 | 88 | 88 | 50 | 88 |

## Discussion

### Model performance

This pre-clinical study demonstrated that HipSAFE models consistently outperformed naïve operators and radiologists without specific training in reading porcine cadavers imaging. It achieved metrics comparable to x-ray imaging with a sensitivity of >90% [25]. HipSAFE’s metrics suggests an opportunity for deep learning in interpretating US imaging collected by operators without sonography training. Mobile-friendly models performed comparably to larger ResNet, demonstrating the viability of edge-computing based approach where the models could be embedded in point of care US devices.

The consistent model performance was likely attributable to the use of high-quality and relatively uniform training data. The feature extractor models were trained on images restricted to key regions of interest, specifically the diaphysis and intertrochanteric regions. In contrast, the target domain consisted of cine clips captured from a broader, unrestricted field surrounding the hip, including muscle, adipose tissue, blood vessels, nerves, connective tissue, and adjacent bones. These additional structures could introduce spurious signals that were not present in the training data.

To address this domain shift, we implemented a simple model-agnostic domain adaptation strategy by adjusting prediction thresholds using the validation operator cine-clip dataset. Although expanding the training dataset to include a wider range of anatomical contexts would be a reasonable alternative, clinical reasoning supports the chosen approach. The target domain is unlikely to introduce new clinically meaningful features indicative of fracture. Instead, previously unseen hyperechoic patterns from surrounding tissues may increase the likelihood of false positives, as intact cortical bone typically appears as a continuous curvilinear signal on ultrasound.

Given that the feature extractors demonstrated high specificity, relatively lower sensitivity, and consistently high AUROC, threshold calibration represented a low-cost and effective adjustment. Aggregating predictions across frames using a simple moving average further stabilized predictions within cine clips by reducing frame-level noise and transient artifacts, thereby improving overall clip-level performance.

### Comparison to other hip fracture detection

To our knowledge, this is the first study evaluating deep learning models for hip fracture detection on US imaging in pre-clinical or clinical populations. Prior work in deep learning hip fracture detection has largely focused on x-rays or computed tomography which have shown similar strong performance [17], [26], [27], [28]. While US is not typically included in standard hip trauma assessment, previous works have established a 95% (95% CI: 94 – 100%) sensitivity and 70% (95% CI: 61 – 79%) specificity on patients [13], [14], [15], which is similar to our pre-clinical deep learning model performance.

Naïve operators underperformed relative to both deep learning models and radiologists, which was anticipated given the technical difficulty of identifying subtle cortical step-offs and inexperience in reading medical imaging. They tend to overcall fractures, which would be a safer approach as a screening tool. Given the lower quality US images and lack of prior experience reading porcine US imaging, the radiologists had reasonable sensitivity and specificity.

### Limitations

The sample size was limited by the cost of conducting a pre-clinical study with porcine cadavers. In comparison to typical hip fracture presentation in older populations, this pre-clinical study had access to juvenile porcine cadavers with open proximal femoral growth plates. The operators and radiologists were blinded to the presence of the open growth plates. Growth plates have similar appearance to cortical bone discontinuities seen in fractures due to greater transmission through the periosteal-cartilage interface than the periosteal-bone interface. Our attribution maps confirm this failure mode as image frames containing intact bone and growth plate are commonly classified as fractured with substantial prediction contributions from the growth plate (and even surpassing the true fracture contributions when it was present). Despite including images with growth plates in the training and validation data, the deep learning models had difficulty learning separatable feature representations between the growth plate and fractures. This anatomical feature resulted in higher false positive detections, suggesting our precision/PPV and specificity may represent a lower bound for clinical studies.

Juvenile femurs have other physiological differences that would affect US images. Their hips do not have osteophytes, which are bony projections as a result of degenerative changes, juveniles have a higher muscle to adipose tissue ratio, and have elastic epimysium and intermuscular fascia. All of these would likely affect model performance because they affect image quality and may introduce spurious features on cine clips.

The performance of the diagnostic radiologists participating in this study does not directly reflect the true performance of detecting hip fractures in a clinical population. The radiologists were trained in interpreting human hip ultrasound images acquired by expert sonographers. Furthermore, they were restricted to unilateral cine clips rather than bilateral comparison, which would decrease diagnostic accuracy.

Naïve operators without sonography or anatomy training were used as a proxy measure for paramedics and nurses’ diagnostic performance. They would likely underestimate the performance achievable by trained healthcare professionals.

Imaging was performed using a single ultrasound transducer and system configuration. The diagnostic performance would be affected by the number and location of the focal region, time gain compensation, imaging depth, and transducer type.

### Considerations for future work

The results suggest that deep learning has the potential to accurately interpret hip US imaging and holds promise for hip fracture detection in human populations. Future work should include larger, clinical datasets, and incorporation a broader range of ultrasound configurations and transducer types. Prospective clinical validation in pre-hospital and emergency settings will be critical to determine its validity and its potential role in improving hip fracture triage and time-to-surgery outcomes.

## Conclusion

In this preclinical study, we evaluated multiple deep learning architectures for automated hip fracture detection on US imaging using porcine cadavers. By assessing both frame-level and cine clip–level predictions, this study demonstrated that both standard and mobile-friendly deep learning models can accurately identify hip fractures. Attribution maps confirmed deep learning model predictions were based on regions within the US images that visualize intact and fractured bone.

While all deep learning models achieved good diagnostic performance, EfficientNet-Lite0 was a lightweight model that achieved the best metrics. Deep learning offers the potential to improve the reliability of interpreting US images by reducing operator dependence, particularly in environments where musculoskeletal US expertise is limited.

Future work should explore the potential clinical role deep learning with hip US as part of a broader hip trauma triaging protocol, especially in rural and resource-constrained settings. Similar approaches to introducing diagnostic imaging to pre-hospital care may contribute to earlier triage, improved diagnostic confidence, and shorten time-to-surgery for patients with suspected hip fractures.

